# Integrating Drug-like Moieties and Binding Site Evolution for Kinase Inhibitor Prediction Using Ensemble Learning Models

**DOI:** 10.1101/2025.05.28.656738

**Authors:** Wei-lin Lin, Yen-Chao Hsu, Jinn-Moon Yang

## Abstract

Protein kinases play a pivotal role in regulating cellular signaling pathways, and their dysregulation is closely associated with numerous diseases, including cancer, autoimmune disorders, and inflammation. Although over 100,000 kinase inhibitors have been developed, only a small fraction has achieved FDA approval, primarily due to off-target effects stemming from the high conservation of kinase binding sites. To address this challenge, we present an ensemble learning framework that integrates both chemical and protein-level information to improve the prediction of selective kinase inhibitors. On the compound side, we construct a 1,048-dimensional feature representation encompassing topological fingerprints, drug-like moieties, atomic composition, and stereochemical descriptors. On the protein side, we develop a 1,700- dimensional representation of kinase binding site environments using multiple sequence alignment and evolutionary conservation information. Comprehensive evaluations across 131 human kinases show that the integration of these features significantly improves model performance, achieving 93.6% accuracy on an independent test set. Furthermore, SHAP-based model interpretation reveals that high-impact features correspond to known binding motifs, such as the P-loop, Hinge region, and DFG motif, as confirmed by crystal structure validation. Lastly, we apply the model to a curated dataset of flavonoid-like compounds, identifying potential natural product-derived kinase inhibitors. This study demonstrates that the proposed integrative approach not only enhances predictive accuracy but also provides interpretable insights into kinase-ligand interactions, offering a promising direction for rational kinase inhibitor design.

## Introduction

Protein kinases play pivotal roles in cellular processes such as cell growth, differentiation, apoptosis, and metabolism, and are thus considered essential drug targets in oncology and inflammatory diseases^1^. Since the approval of the first kinase inhibitor, imatinib, for chronic myeloid leukemia, more than 70 kinase-targeted drugs have been approved by the U.S. FDA, with hundreds more under clinical evaluation^2^.

Despite this success, a significant challenge remains in the selective targeting of kinases. Many kinases share highly conserved ATP-binding pockets, making off-target interactions and resistance mutations common issues^3^. To overcome these problems, various kinase classification systems have been developed to better understand kinase diversity and guide inhibitor design^4^.

Machine learning (ML)-based approaches have shown great promise in predicting kinase–ligand interactions by learning patterns from biochemical datasets^5^. Among these, proteochemometric modeling (PCM), which integrates both ligand and protein features, has demonstrated improved performance in multi-target settings^6^. However, the representation of protein features in such models remains an open issue. Commonly used sequence encoders such as ESM or ProtT5 often produce high-dimensional embeddings that lack interpretability, and typically encode the entire protein rather than the functionally relevant binding regions.

In this study, we introduce a pairwise kinase–compound prediction framework that leverages interpretable feature design for both ligands and kinases (Figure 1). For protein descriptors, we constructed a curated multiple sequence alignment (MSA) across 419 human kinases, focusing on 85 conserved binding site residues to capture structural and evolutionary information^7^. For compound representation, we extracted 1,048 interpretable chemical features, including ECFP fingerprints, substructure patterns, ring systems, and stereochemical markers, enabling SHAP-based interpretability analysis.

**Figure 1.**
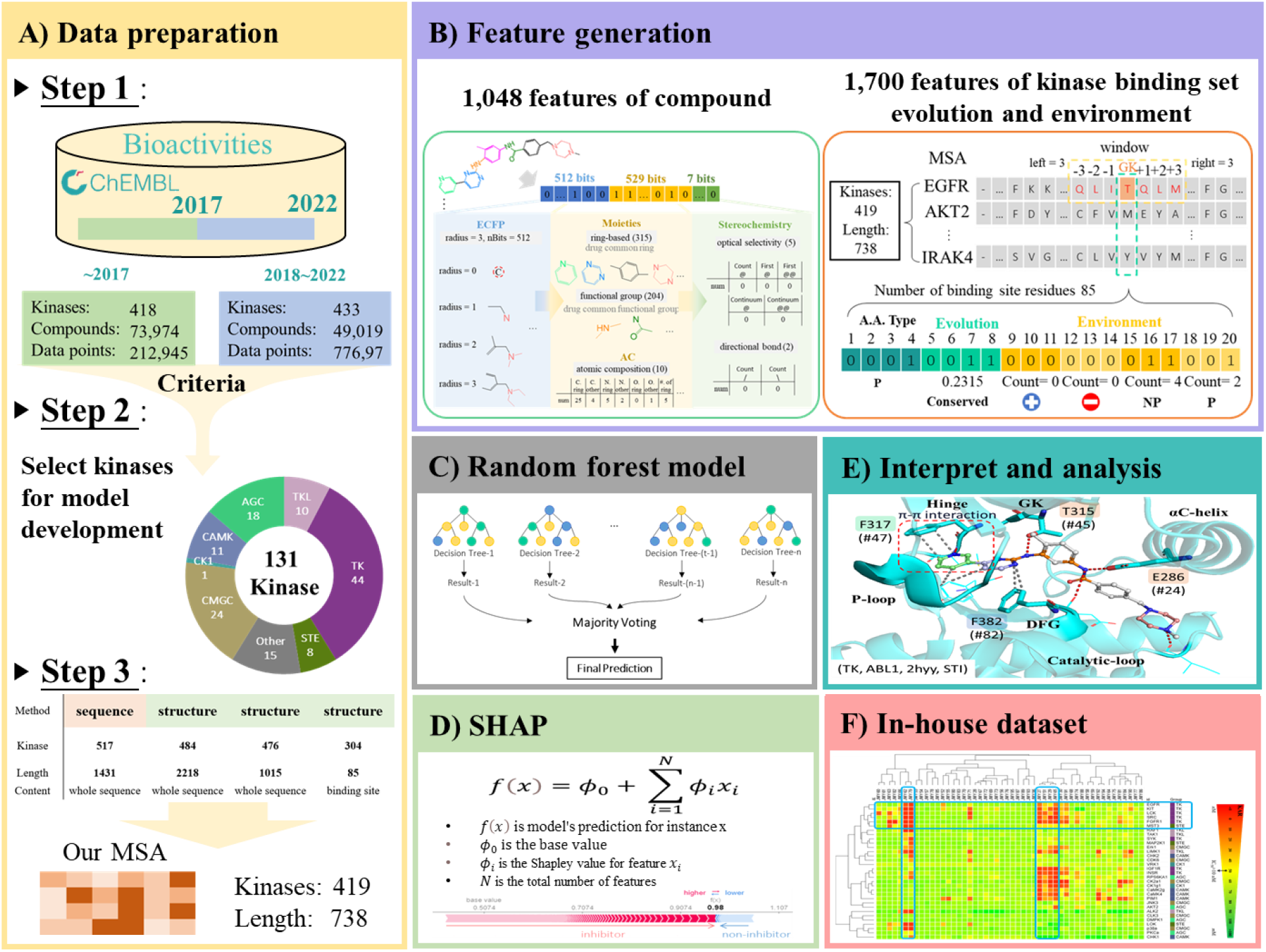
Overview of the integrative prediction framework for kinase inhibitor selectivity. (A) Data preparation: Kinase–inhibitor interaction data were collected from ChEMBL and time-split into a training/test set (pre-2017, 80:20) and an independent validation set (2018–2022), covering 131 kinases. A custom multiple sequence alignment (MSA) of 419 kinases (sequence length: 738 residues) was generated using both sequence and structure-based references. (B) Feature generation: Each compound was encoded using 1,048 binary features based on drug-like substructures, ring systems, and stereochemistry. Each kinase was represented using 1,700 features derived from residue-level evolutionary conservation and local binding site environments within the MSA. (C) Random Forest models were trained separately for each kinase using ensemble learning. (D) SHAP (Shapley Additive Explanations) was applied to quantify the contribution of each feature to prediction outcomes. (E) Structural interpretation: High-impact protein features were mapped onto kinase crystal structures to validate their biochemical relevance (e.g., hinge region, P-loop, DFG motif). (F) In-house dataset application: The model was applied to a curated collection of flavonoid-like compounds to evaluate their potential as multi-target kinase inhibitors.

Our model was trained using Random Forest classifiers on time-split data from ChEMBL, separating pre-2017 data for training and post-2018 data for independent evaluation^8^. We applied the model to an in-house flavonoid-like compound dataset to investigate potential multi-target kinase inhibition and to validate biochemical relevance via SHAP-derived feature interpretation.

## Materials and Methods

### Dataset Collection and Preprocessing

We constructed a kinase–ligand interaction dataset by integrating and filtering bioactivity data from ChEMBL version 23 and version 31. To ensure a realistic evaluation of model generalizability, we performed a time-split strategy: all data published before 2017 were used for training and testing (80:20 split), and those from 2018 to 2022 were reserved as an independent test set.

Bioactivity records were filtered based on the following criteria:

1. Target organism: Homo sapiens protein kinases (as defined by UniProt).
2. Target type: single protein.
3. Assay type: binding.
4. Bioactivity types: IC_50_, EC_50_, K_i_, K_d_, or AC_50_ with valid pChEMBL values.
5. Duplicate records were collapsed using median bioactivity.

We transform them into binary labels based on the following definition: when the measured biological activity value of compound *C*_*i*_ in target kinase *K* is ≤ 10 μM, it is defined as inhibitor (Label: 1); when the measured biological activity value of compound *C*_*i*_ in target kinase *K* is ≥ 20 μM, it is defined as non-inhibitor (Label: 0), as shown in the formula below:

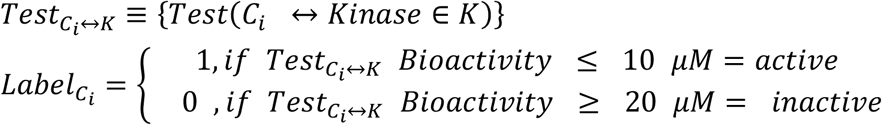

After classification, the dataset prior to 2017 includes 417 protein kinases and 72,837 compounds, with 108,967 active inhibitors and 101,228 inactive inhibitors (before expansion, there were only 4,382 entries in the inactive inhibitors category). For the period 2018 to 2022, there are 429 protein kinases and 46,372 compounds, comprising 72,134 active inhibitors and 2,001 inactive inhibitors (Figure 2).

**Figure 2.**
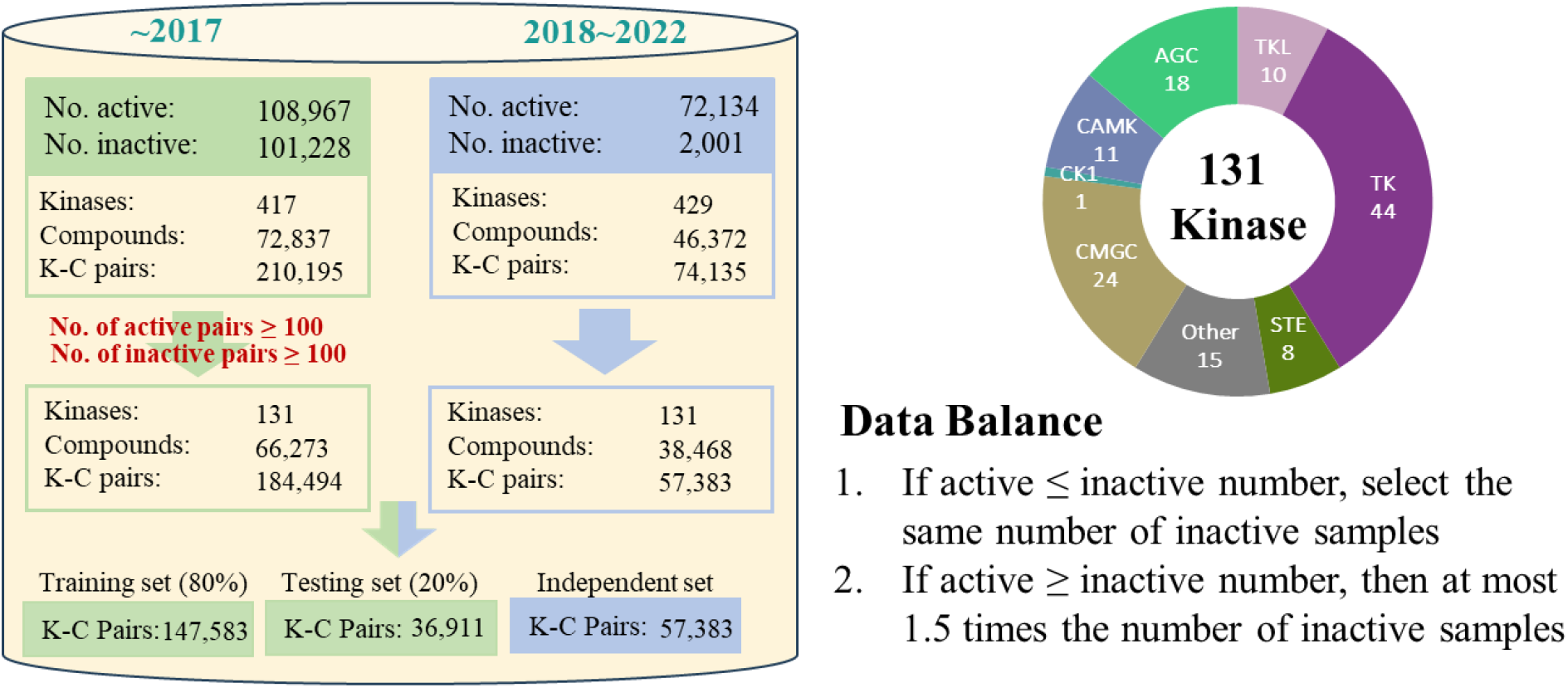
Screening kinases and data sets to build a model.

### Compound Feature Engineering

Each compound was encoded using a 1,048-dimensional feature vector (Figure 3) composed of:

**Figure 3.**
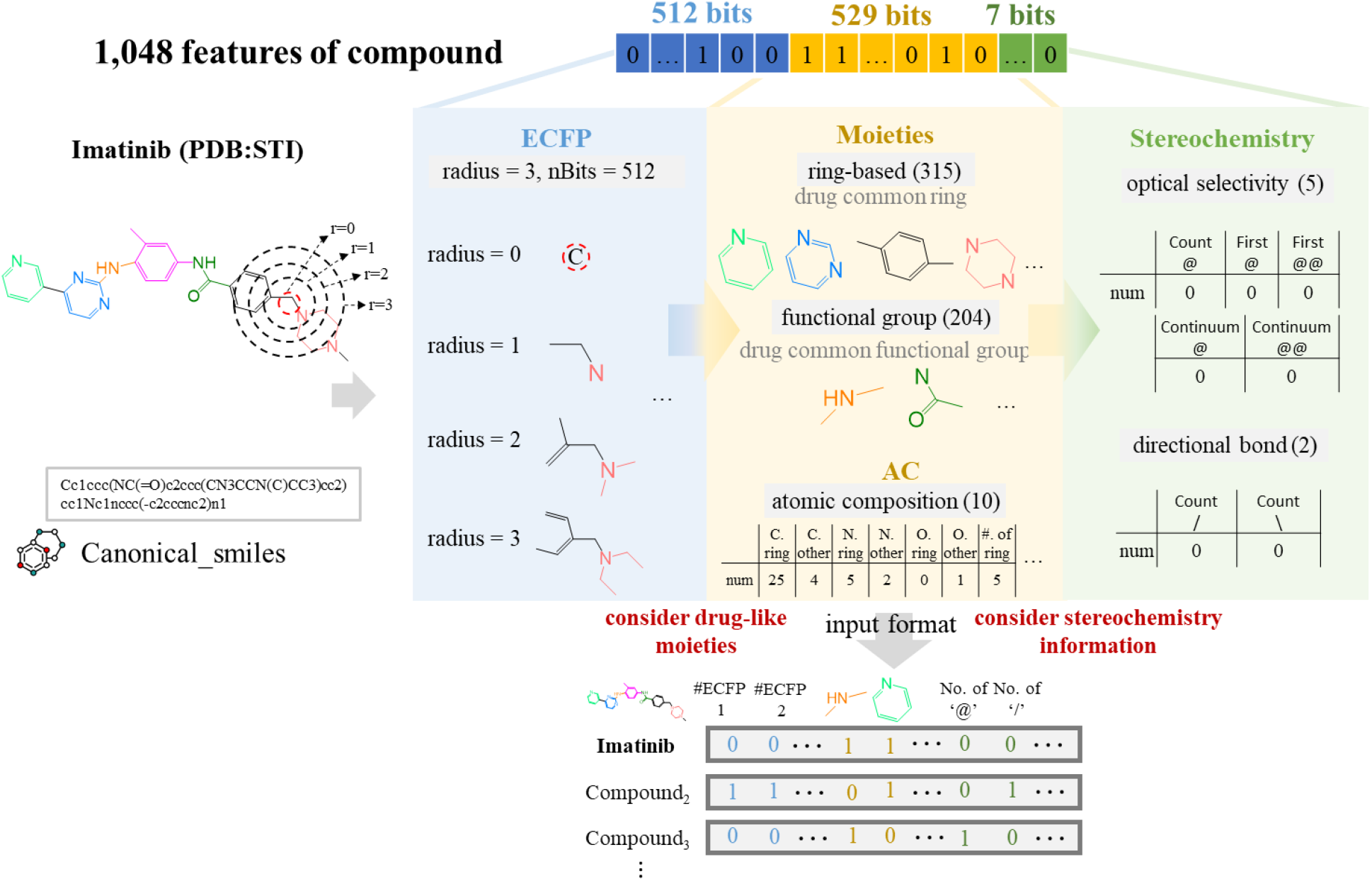
Construction of 1,048 compound features incorporating drug-like moieties and stereochemical information.

1. Topological fingerprints (512 features): Extended-Connectivity Fingerprints (ECFP4), generated via RDKit.
2. Functional group descriptors (204 features): Presence of common substructures defined in Checkmol.
3. Ring system descriptors (315 features): Extracted from PubChem Ring and Ring-in-Drug datasets.
4. Atomic composition (10 features): Counts of selected atoms and ring structures.
5. Stereochemical descriptors (7 features): Including chiral center notations (@, @@), cis/trans markers (“/”, ““) and their positions or counts.

All compound features were converted into binary or normalized numerical format.

### Protein Feature Engineering

We constructed multiple sequence alignments (MSA) for 419 human kinases based on sequence and structure-based methods (Table 1, including alignments from Creixell et al.^7^, Gustafsson et al.^9^, and van Linden et al.^10^). To minimize gaps and retain pocket-centric alignment, we manually curated binding site regions and filtered out degenerate or pseudokinase domains.

**Table 1.**
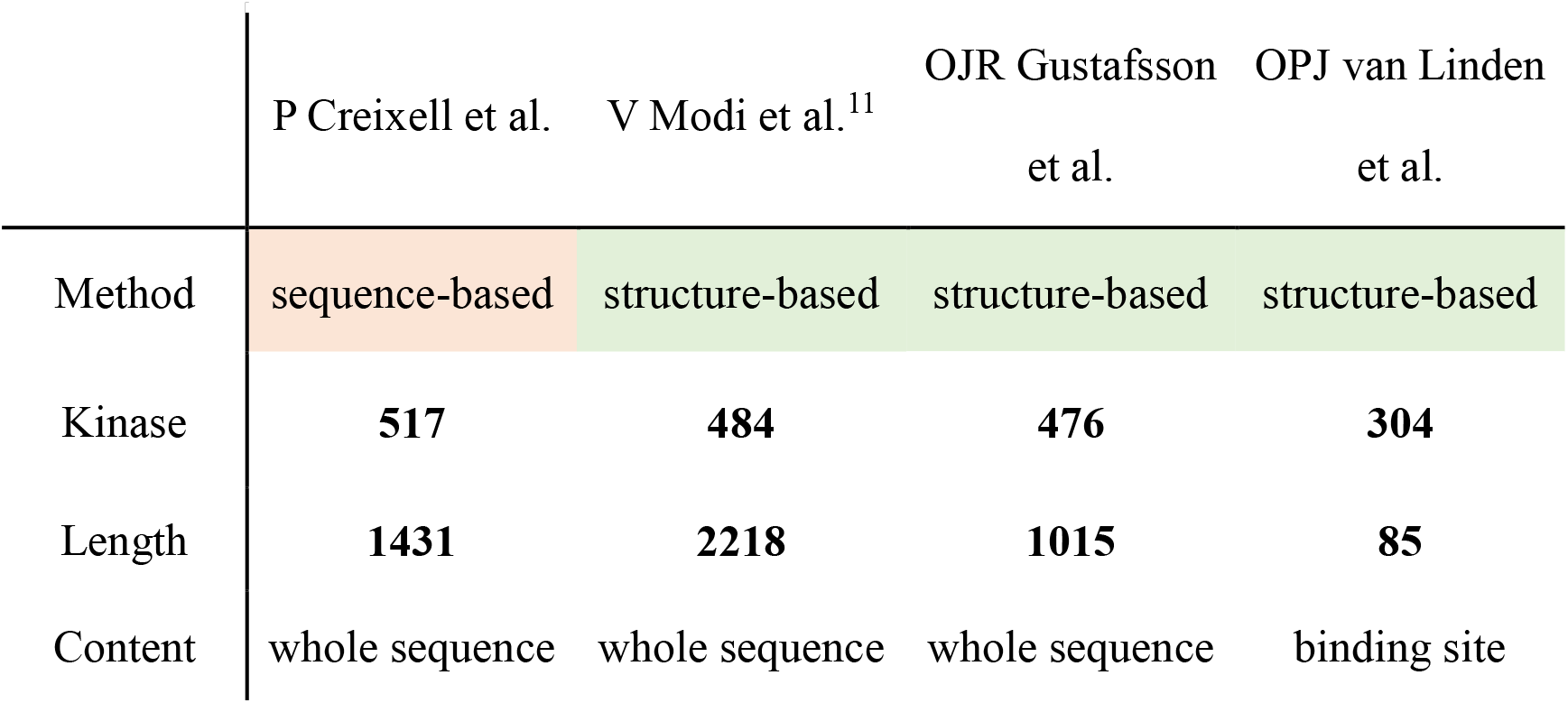
Collection sequence-based and structure-based MSA.

|  | P Creixell et al. | V Modi et al.<sup>11</sup> | OJR Gustafsson et al. | OPJ van Linden et al. |
| --- | --- | --- | --- | --- |
| Method | sequence-based | structure-based | structure-based | structure-based |
| Kinase | **517** | **484** | **476** | **304** |
| Length | **1431** | **2218** | **1015** | **85** |
| Content | whole sequence | whole sequence | whole sequence | binding site |

Each kinase was encoded as a 1,700-dimensional vector derived from 85 binding site residues (Figure 4). For each residue, we extracted:

**Figure 4.**
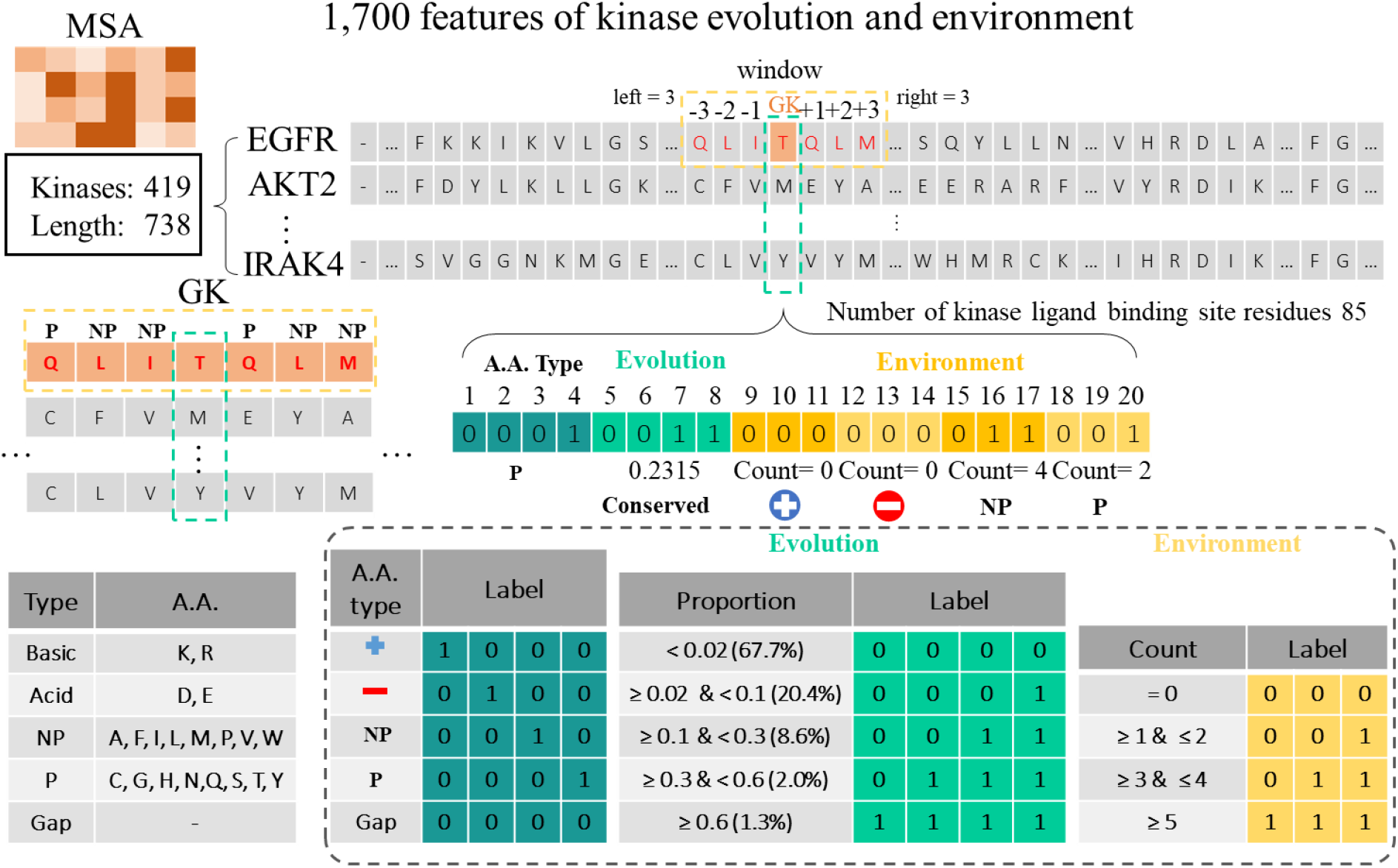
Protein feature descriptors encoding flowchart. Construction of 1,700 kinase binding site descriptors from multiple sequence alignment.

1. Amino acid type (4 bits): Categorized into basic, acidic, non-polar, or polar groups.

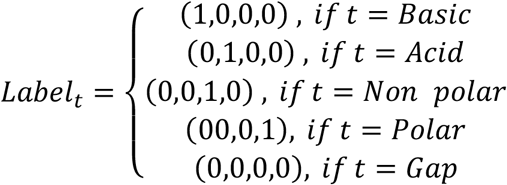
2. Conservation (4 bits): Discretized based on residue frequency at aligned position across kinases.

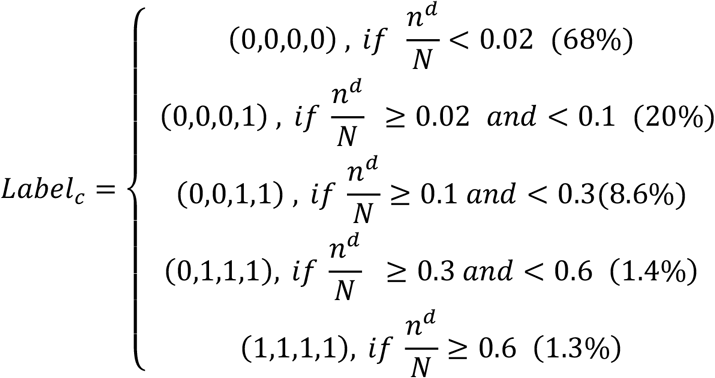
3. Local environment (12 bits): One-hot representation of surrounding amino acid categories within a ±3 residue window.

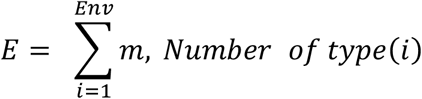

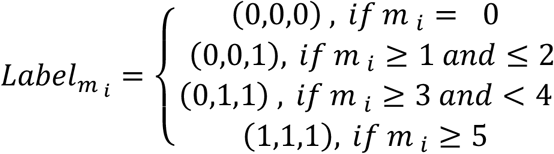

This static kinase representation was assigned uniformly across all pairs involving the same protein.

### Model Architecture

We employed a Random Forest (RF) classifier for each kinase using the Scikit-learn implementation. Each model was trained independently per kinase using the selected active/inactive compound pairs. Hyperparameters were set as follows:

1. Number of trees: 500
2. Max depth: None
3. Feature selection: sqrt(# features) per split
4. Class balancing: stratified sampling

Performance was evaluated on held-out test sets and the independent post-2017 dataset.

### Feature Importance and Model Interpretation

We computed feature importance using two complementary approaches:

1. Gini-based importance from Random Forest splitting criteria.
2. SHAP (Shapley Additive Explanations) values to assess contribution directionality and contextual impact on predictions.

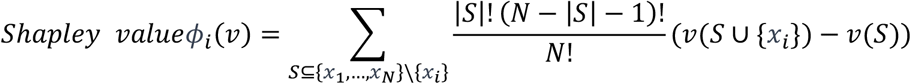

SHAP values were averaged over 500 test samples per kinase and visualized to identify consistent high-impact features.

### Flavonoid Case Study

To explore real-world applicability, we assembled an in-house dataset of 50 flavonoid-like compounds tested against 32 kinases. These molecules shared common scaffolds with ATP and known kinase inhibitors. The same encoding and prediction pipeline were applied to evaluate their potential activity and selectivity profiles.

## Results and Discussion

### Compound Feature Integration Enhances Prediction Performance

We first evaluated the impact of compound feature engineering on kinase inhibitor prediction. Using only compound features, we trained Random Forest classifiers for 131 kinases. Compared to models using standard ECFP fingerprints alone, the incorporation of drug-like moieties—such as functional groups, ring systems, atomic composition, and stereochemical descriptors—led to performance improvements in 116 kinases. (Figure 5)

**Figure 5.**
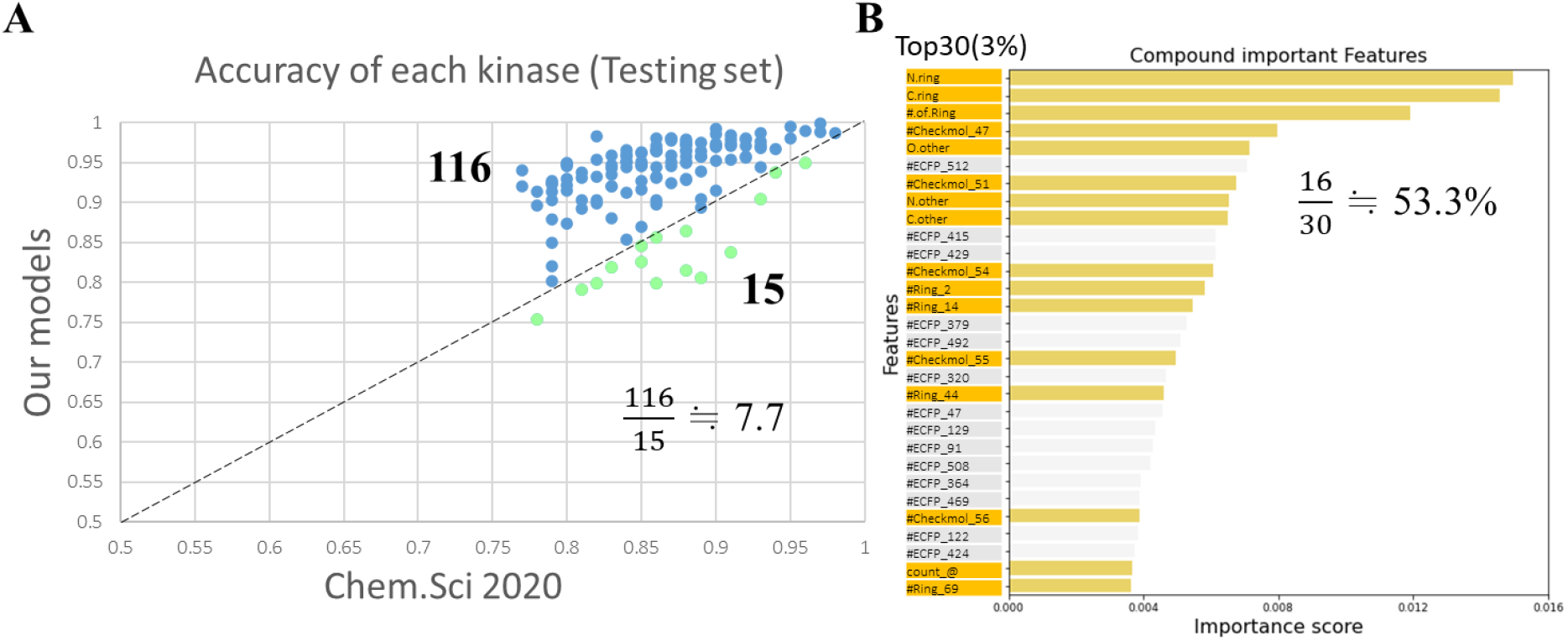
Comparison of model performance and importance feature ranking.

Among these, kinases with more structurally diverse ligands, such as EGFR, BTK, and CDK2, exhibited the most significant gains. These results suggest that the inclusion of chemically interpretable substructures expands the model’s understanding of ligand selectivity.

Feature importance analysis revealed that over 50% of the top 3% most informative features originated from the newly added moiety-based descriptors rather than traditional fingerprints. This highlights the value of incorporating domain knowledge into compound encoding for interpretability and performance.

### Incorporating Binding Site Evolution Improves Generalization

To further enhance model robustness and kinase-specific learning, we integrated protein binding site features derived from multiple sequence alignments. With the 1,700-dimensional kinase representations added, all 131 kinase models demonstrated improved predictive performance on both internal and independent datasets.

Notably, the average accuracy on the independent time-split dataset increased from 87.4% (compound-only) to 93.6% (compound + protein features). Kinases with high sequence similarity but divergent ligand preferences—such as JAK1 vs. JAK3— benefited significantly from the addition of evolutionary context, confirming that residue-level conservation and local biochemical environment are key determinants of inhibitor specificity.

### SHAP-Based Interpretation Reveals Mechanistic Consistency

We applied SHAP (Shapley Additive Explanations) to decompose individual predictions and identify key features contributing to kinase inhibition. Across multiple kinases, the most impactful compound features often corresponded to known bioactive substructures (e.g., heteroaromatic rings, hydrogen bond donors).

On the protein side, SHAP analysis identified several conserved regions— including the P-loop, hinge region, and DFG motif—as frequent contributors to positive predictions. These residues are well-established as critical for ATP or ligand binding. We validated these findings by mapping SHAP-highlighted residues onto crystal structures (e.g., ABL1–imatinib complex), confirming spatial overlap with binding pockets. These results demonstrate that the model not only achieves high predictive accuracy but also captures biologically meaningful interaction patterns.

### Kinome-Wide Evaluation and Group-Level Patterns

Using hierarchical clustering of kinase features and model outputs, we observed that predictive performance generally aligned with known kinase families in the human kinome. (Table 2) For instance, CMGC kinases (e.g., CDKs, GSK3, MAPKs) showed consistent predictive performance due to well-characterized ligand spaces, while atypical and less-characterized kinases exhibited greater variance.

**Table 2.** Performance differences of kinase models on different datasets.

| Kinase | Group | (Independent set) |  |  | (Testing set) |  |  | Acc<sub>i</sub>-Acc<sub>t</sub> |
| --- | --- | --- | --- | --- | --- | --- | --- | --- |
|  |  | act | inact | Acc<sub>i</sub> | act | inact | Acc<sub>t</sub> |  |
| COT | STE | 25 | 0 | **0.0%** | 33 | 33 | **98.5%** | **-98.5%** |
| PBK | Other | 40 | 0 | **0.0%** | 21 | 21 | **97.6%** | **-97.6%** |
| FGFR3 | TK | 898 | 0 | **4.5%** | 107 | 107 | **94.4%** | **-89.9%** |
| CDK6 | CMGC | 72 | 3 | **10.7%** | 57 | 57 | **100.0%** | **-89.3%** |
| IRE1 | Other | 58 | 11 | **7.2%** | 60 | 60 | **95.8%** | **-88.6%** |
| DLK | TKL | 9 | 0 | **100.0%** | 130 | 130 | **98.8%** | **1.2%** |
| PKCh | AGC | 12 | 0 | **100.0%** | 42 | 42 | **98.8%** | **1.2%** |
| CHK2 | CAMK | 228 | 2 | **93.5%** | 53 | 53 | **91.5%** | **2.0%** |
| PYK2 | TK | 46 | 4 | **82.0%** | 39 | 39 | **76.9%** | **5.1%** |

We also observed that kinases with higher evolutionary entropy in the binding site region tended to rely more heavily on compound features, whereas those with lower entropy (i.e., conserved pockets) benefited more from protein features. This suggests a functional link between binding site conservation and feature modality importance.

### Application to Flavonoid-Like Compounds

To evaluate the model’s practical utility, we applied it to a curated panel of 50 flavonoid-like compounds tested against 32 kinases (Figure 6). Despite the structural simplicity of flavonoids, the model successfully identified several compound–kinase pairs with high predicted binding probability, including interactions with PIM1, CK2α1, and FGFR1.

**Figure 6.**
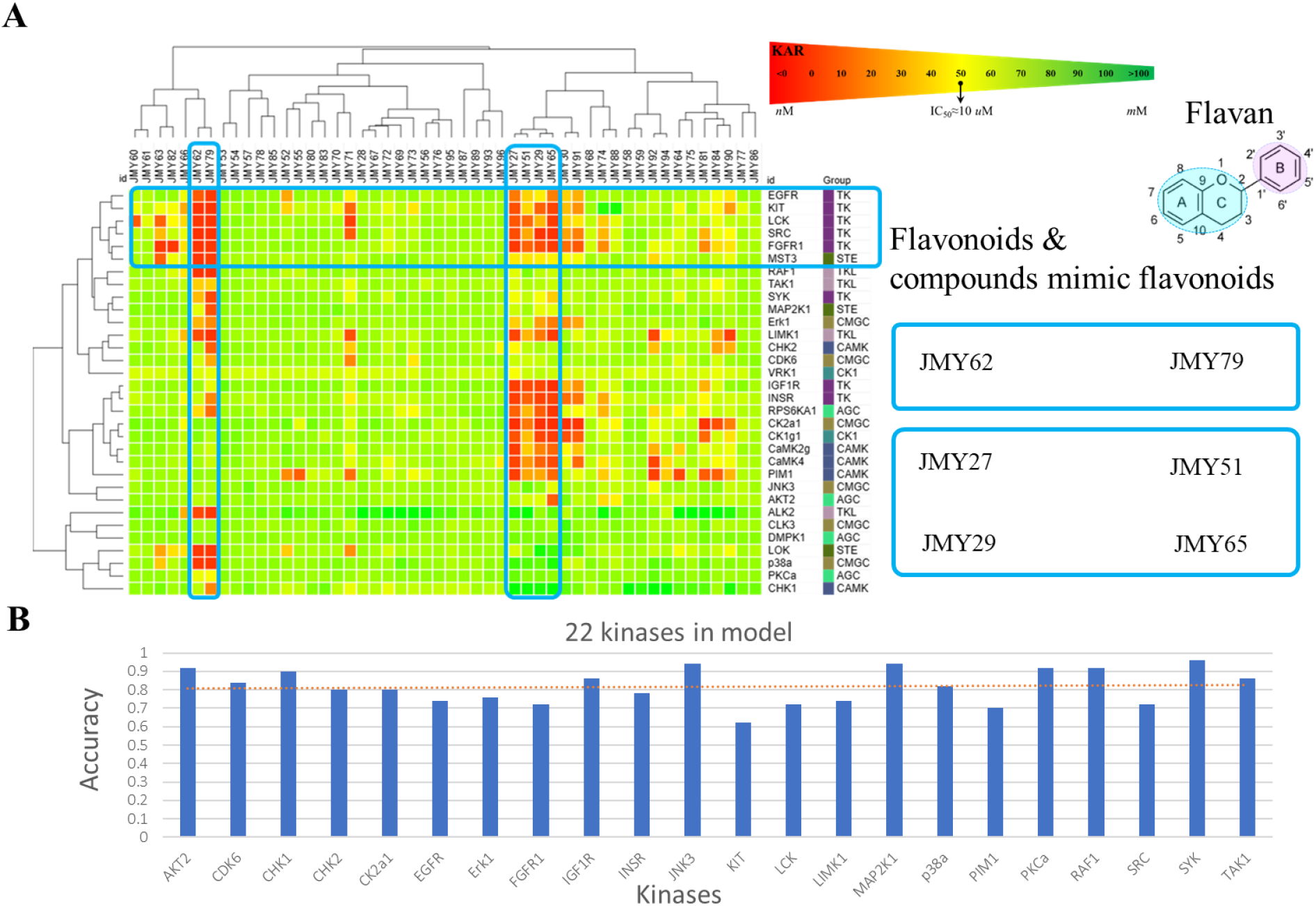
Analysis and application of flavonoid-like kinase profiling.

Interestingly, SHAP analysis of these predictions revealed that hydroxylated aromatic rings—common in flavonoids—were recognized as key activating substructures, and the corresponding kinase pockets often featured polar or charged residues in the hinge region. This aligns with literature reports of flavonoids as moderate multi-target kinase inhibitors, and demonstrates the model’s capacity to generalize to natural products with novel scaffolds.

### Discussion: Implications and Limitations

Our results underscore the importance of integrating chemically interpretable features with biologically meaningful protein representations to enhance both performance and explainability in kinase inhibitor prediction.

However, one limitation of the current model is that protein features remain fixed across all compound pairs for a given kinase. While this design is computationally efficient and leverages conserved evolutionary information, it may restrict the model’s ability to capture pair-specific binding conformations or ligand-induced pocket flexibility. Future efforts may incorporate pair-aware features, such as ligand-conditioned binding site embeddings or docking-based contact maps, to more accurately reflect interaction specificity.

Additionally, although Random Forests offer high interpretability, they may underperform compared to deep learning methods on large-scale datasets. Expanding this framework to incorporate graph neural networks or attention-based architectures may further improve performance, especially for low-data or unseen targets.

## Conclusion

In this study, we developed an interpretable and scalable machine learning framework for kinase inhibitor prediction by integrating compound-level moiety descriptors and protein-level binding site evolution features. Through comprehensive evaluation across 131 human kinases, we demonstrated that this dual-feature approach significantly improves predictive performance and generalizability compared to traditional compound-only models.

Our feature engineering strategy extends beyond conventional molecular fingerprints by incorporating chemically meaningful substructures, stereochemistry, and atomic composition—features that contribute directly to ligand selectivity. On the protein side, the inclusion of residue-level evolutionary conservation and local context enables the model to better capture the structural and biochemical determinants of kinase specificity.

Importantly, the use of SHAP-based interpretation reveals that our model not only achieves high accuracy but also aligns with known interaction motifs in kinase– ligand complexes, offering valuable mechanistic insights. The successful application of our model to flavonoid-like natural compounds further illustrates its potential to guide the discovery of novel inhibitors from diverse chemical spaces.

Overall, this work contributes a biologically grounded, chemically informed, and computationally efficient strategy for kinase inhibitor prediction. Our integrative approach bridges the gap between ligand-centered and protein-centered models, and provides a robust platform for rational drug design, selectivity profiling, and kinase target expansion.

